# A computational pipeline to predict cardiotoxicity: From the atom to the rhythm

**DOI:** 10.1101/635433

**Authors:** Pei-Chi Yang, Kevin R. DeMarco, Parya Aghasafari, Mao-Tsuen Jeng, Sergei Y. Noskov, Vladimir Yarov-Yarovoy, Igor Vorobyov, Colleen E. Clancy

## Abstract

We simulate and predict cardiotoxicity over multiple temporal and spatial scales from the drug chemistry to the cardiac rhythm.

**ABSTRACT:** Drug-induced proarrhythmia is so tightly associated with prolongation of the QT interval that QT prolongation has become widely accepted as a surrogate marker for arrhythmia. The problem is that QT interval as an arrhythmia indicator is too sensitive and not selective, resulting in many potentially useful drugs eliminated early in the drug discovery process. We first set out to predict the fundamental mode of binding for the proarrhythmic drug dofetilide with the promiscuous cardiac drug target, the hERG potassium channel. In a novel linkage between the atomistic and functional scales, computed binding affinities and rates from atomistic simulation are utilized here to parameterize function scale kinetic models of dofetilide interactions with the hERG channel. The kinetic model components are then integrated into predictive models at the cell and tissue scales to expose fundamental arrhythmia vulnerability mechanisms and complex interactions underlying emergent behaviors. Human clinical data from published studies were used to validate model framework and showed excellent agreement, demonstrating feasibility of the approach. The model predictions show that a clinically relevant dose of dofetilide increased arrhythmia vulnerability in all emergent TRIaD-linked parameters including Triangulation, Reverse use-dependence, beat-to-beat Instability and temporal and spatial action potential duration Dispersion. Application of machine learning demonstrated redundancy in the TRIaD linked parameters and suggested that changes in beat-to-beat instability were highly predictive of arrhythmia vulnerability in this setting. Here, we demonstrate the development and validation of a prototype multiscale model framework to predict electro-toxicity in the heart for the proarrhythmic drug dofetilide from the atom to the rhythm.

**SIGNIFICANCE STATEMENT:** Cardiotoxicity in the form of deadly abnormal rhythms is one of the most common and dangerous risks for drugs in development and clinical use. There is an urgent need for new approaches to screen and predict the effects of chemically similar drugs on the cardiac rhythm ***and*** to move beyond the QT interval as a diagnostic indicator for arrhythmia. To this end, we present a computational pipeline to predict cardiotoxicity over multiple temporal and spatial scales from the drug chemistry to the cardiac rhythm. We utilize predicted quantitative estimates of ion channel-drug interactions from our companion paper to simulate cardiotoxicity over multiple temporal and spatial scales from the drug chemistry to the cardiac rhythm.

## INTRODUCTION

Cardiotoxicity is one of the most common reasons for drug removal from the market, typically manifesting as prolongation of the QT interval on the ECG with the potential for fatal ventricular arrhythmias (1). In the context of drug induced cardiac arrhythmia, the vital hindrance to prevention of electrical rhythm disturbances is a lack of meaningful approaches to ***distinguish between therapeutic, benign or harmful*** actions of drugs. An important example is the use of QT interval prolongation as a surrogate marker for proarrhythmia (2). Since 2005, the regulatory process for preclinical drug candidates includes a dedicated clinical study in healthy volunteers, the so-called “Thorough QT Study” (TQT) (3). A drug that results in greater than 5 ms QT prolongation above normal in healthy humans indicates “regulatory concern” (4).

The problem is that many useful drugs that were approved prior to the 2005, including the commonly used ***antiarrhythmic agents*** verapamil, ranolazine and amiodarone to name a few, all fail the TQT – they all block K+-selective cardiac channel hERG and prolong QT interval (5–7). Indeed, there exist numerous examples of safe and effective drugs (including antiarrhythmics, antipsychotics and antibiotics) that gained FDA approval prior to TQT implementation (1, 2, 8). If screened today, these safe drugs would fail the test.

Abnormal cardiac electrical activity is most often a side effect from unintended block of the promiscuous drug target hERG, the delayed rectifier K+ channel in the heart. hERG block results in prolongation of the QT interval on the ECG, a phase of the cardiac cycle that corresponds to ventricular repolarization.

Not all hERG block is proarrhythmic. But, at present, there is no accurate method to distinguish unsafe hERG blockers from safe drugs. There are at least two distinct classes of hERG blockers that prolong QT interval (1): In the first group are drugs that block hERG, prolong QT interval and increase proclivity to potentially deadly torsades de pointes (TdP) arrhythmias. This group includes, for instance, sotalol, dofetilide, and cisapride (9, 10). The second group consists of hERG blockers that prolong QT interval and have low risk for ventricular arrhythmias like ketaconozole, moxifloxacin, amiodarone, and verapamil (11–16).

There have been many attempts to distinguish the two classes of hERG blockers via “top-down” approaches including studies devoted to analytical methods aimed at assessing the relationship between rate corrected QT “morphology” and arrhythmia risk (17–20). More recently, the Comprehensive in Vitro Proarrhythmia Assay (CiPA) initiative comprising a worldwide team of regulators, academicians and industry scientists have worked to develop and validate a collection of nonclinical assays to move beyond the thorough-QT study and better predict preclinical arrhythmia risk (21). This program promotes screening of additional cardiac ion channel targets and assessing net impact on the cellular action potential duration and QT interval via *in vitro* experiments and *in silico* functional models. While the consideration of multi-channel block is an improvement, even these newer computational models still generally rely on IC50 values as model inputs, exclude non-ionic drug targets and rely on prolongation of the action potential duration or QT interval as the marker for safety (22–26). Moreover, although these methods have shown to be fairly effective at categorizing risk for the CiPA test compounds, there has not yet been mechanistic validation of the accuracy of the model prediction. In other words, a test drug may land in the right category, but for the wrong reason. The *in silico* assay currently cannot distinguish between chemically similar drugs with similar affinity profiles. Moreover, incorporating individual genetic differences into the current model schema is not yet possible. Finally, there are a variety of drugs that do not appear to exhibit multichannel block, but rather hERG block alone with low arrhythmia risk such as moxifloxacin, ketaconozole and the selective-serotonin reuptake inhibitor CONA-437 (14, 27).

We hypothesized that channel conformation state specificity and the associated kinetics of hERG block may promote TdP as indicated by the TRIaD: *Triangulation, reverse* use dependence, beat-to-beat *instability* of action potential duration, temporal *and* spatial action potential duration *dispersion (9, 28, 29).* Here, we use an integrative experimental and computational modeling and machine learning approach that spans scales from the atom to the cardiac rhythm to predict the intrinsic properties of the structure-activity relationship that determine proarrhythmia for the prototype drug dofetilide, which is a potent hERG blocker, prolongs QT interval and has a high risk for TdP arrhythmias (30, 31). Thus, it was chosen as an ideal candidate for our proof-of-the-concept study.

In this study, we take the first necessary steps to answer the question, *“Can we predict when a QT prolonging drug is a proarrhythmic drug based its chemistry?”* We have assembled a proof of concept multiscale model for predictive cardiac pharmacology. We present a novel multiscale approach based on structural and dynamic atomistic models of drug-channel interactions and kinetics intended to predict drug-induced arrhythmia. We expect that the model framework may be expanded to make an impact in drug discovery, drug safety screening for a variety of compounds and targets, and in a variety of regulatory processes. We propose that the potential added value in this method is that it can be expanded to incorporate a variety of additional drug targets to glean binding affinities at the atomic scale, and the structural models can be extended to include genetic mutations, polymorphisms or variants of unknown significance that may impact drug binding. The method can be also used to distinguish between chemically similar drugs that may appear to have similar effects on the action potential and QT interval.

## Materials and methods

### Structural modeling of hERG

The wild-type drug-free atomistic structural hERG model, shown in Fig. 1A, was developed via an iterative refinement of a recent cryo-EM structure of hERG (PDB ID: 5VA1) as described in (REF). This model was shown to represent an open conducting state of hERG.

**Fig. 1.**
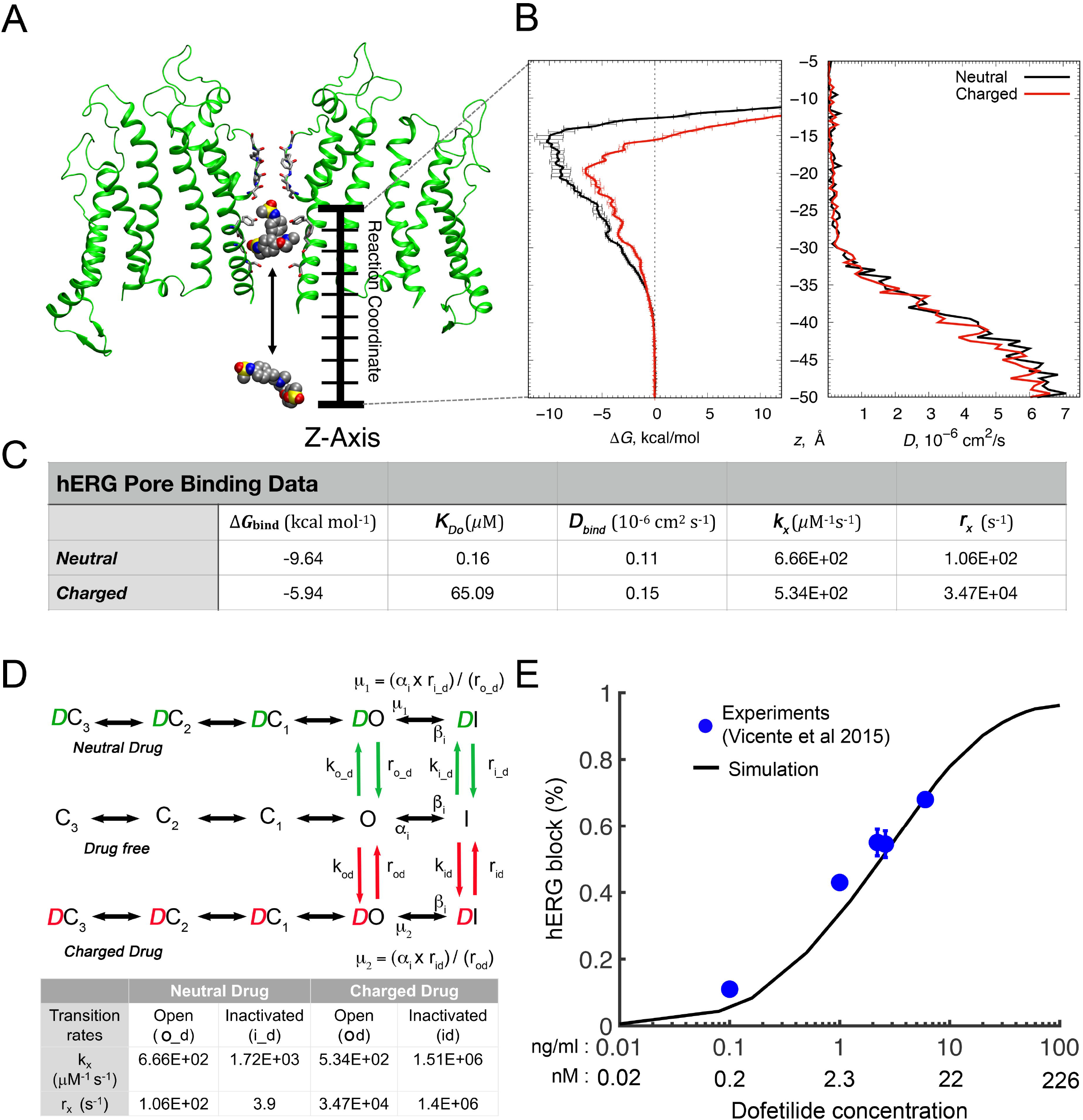
Concentration and open state-dependent block of hERG by dofetilide. (A) A schematic of a molecular system used to compute potential of mean force (PMF) profiles for dofetilide – hERG interactions. hERG structure is shown as green ribbons, dofetilide molecules inside and outside channel pore are shown in space-filling representations. (B) PMF (lef) and diffusion coefficient (right) profiles for open hERG – dofetilide interactions. Dominant low free energy wells of −9.6 and −5.9 kcal/mol at z ≈ – 15 Å and −20 Å were identified for neutral (black) and charged (red) dofetilide in the hERG pore (left). Diffusion coefficients for entry of dofetilide into the open pore showing that charged and neutral drug values are comparable during the binding process (right). (C) Binding affinities and drug diffusion rates used to calculate the drug “on” (*k_x_*) and “off” (*r_x_*) model transition rates for (D) the Markov model open state parameters. The model represents a map of the hERG channel functional states. Drug free (black), charged drug bound (red) states, and neutral drug bound (green) states are shown. Model parameters *K*_DI_ for drug open-inactivated states were reduced 70-fold relative to the open state based on measurements (35). (E) Experimentally measured dose dependent inhibition of hERG by dofetilide (blue circles) (33) and optimized model-based results (black curve).

### Dofetilide-hERG interaction function scale model

The wild-type drug-free hERG Markov model previously described in (32) is shown in Figure 1D. To simulate drug interactions with hERG, we used ***simulated affinities***, (i.e. drug dissociation constants) *K*_D_, and drug diffusion rates, *D*, both computed from the umbrella sampling (US) molecular dynamics (MD) simulations used to constrain the drug “on” (*k_o_d_* and *k_od_*) and “off” (*r_o_d_* and *r_od_*) model transition rates for open state (Figure 1D – bottom). There are two modes of drug bound channel states – neutral (green) and charged (red). The charged and neutral drug fractions, *f*_1_ and *f*_0_, are calculated using the following equations:

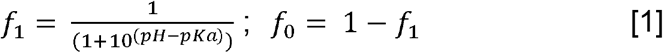

Where *pH* = 7.2 and *pK_a_* = 7.0 (33, 34).

**Table 1:** Transition rates in the I_Kr_ model

| Transition rates ($ms^{-1}$) | |
| --- | --- |
| *Drug free Kr channel* |  |
| C3→C2 | $ae = \frac{T}{T_{base}} e^{(24.335 + \frac{T_{base}}{T}(0.0112 \times V - 25.914))}$ |
| C2→C3 | $be = \frac{T}{T_{base}} e^{(13.688 + \frac{T_{base}}{T}(-0.0603 \times V - 15.707))}$ |
| C2→C1 | $ain = \frac{T}{T_{base}} e^{(22.746 + \frac{T_{base}}{T}(-25.914))}$ |
| C1→C2 | $bin = \frac{T}{T_{base}} e^{(13.193 + \frac{T_{base}}{T}(-15.707))}$ |
| C1→O | $aa = \frac{T}{T_{base}} e^{(22.098 + \frac{T_{base}}{T}(0.0365 \times V - 25.914))}$ |
| O→C1 | $bb = \frac{T}{T_{base}} e^{(7.313 + \frac{T_{base}}{T}(-0.0399 \times V - 15.707))}$ |
| O→I | $\beta i = \frac{T}{T_{base}} e^{(30.016 + \frac{T_{base}}{T}(0.0223 \times V - 30.88))} \times \left(\frac{5.4}{[K]^o}\right)^{0.4}$ |
| I→O | $\alpha i = \frac{T}{T_{base}} e^{(30.061 + \frac{T_{base}}{T}(-0.0312 \times V - 33.243))}$ |

**Table 2:**
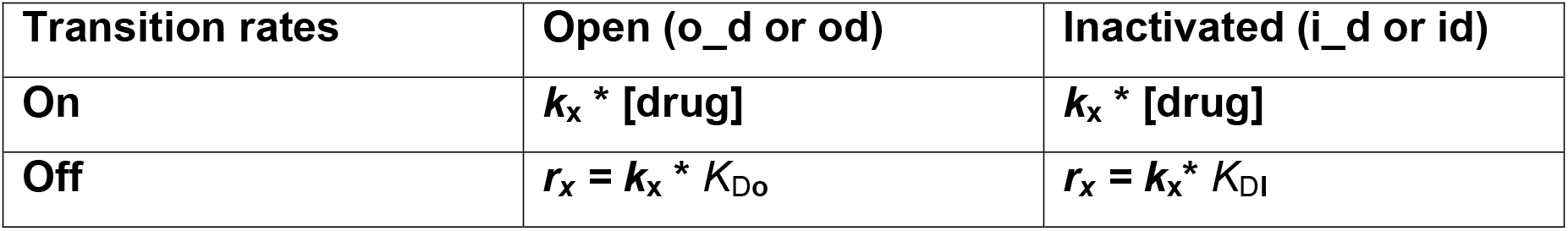
Transition rates for Dofetilide model (see Figure 1D)

| Transition rates | Open (o_d or od) | Inactivated (i_d or id) |
| --- | --- | --- |
| On | $k_x * [\text{drug}]$ | $k_x * [\text{drug}]$ |
| Off | $r_x = k_x * K_{Do}$ | $r_x = k_x * K_{DI}$ |

Dissociation constants (*K*_Do_) as well as *on* and *off* rates of dofetilide for the open state hERG model for neutral and charged drugs are computed from the US MD derived potential of mean force (PMF) i.e. free energy, and diffusion coefficient profiles (Figure 1B). Based on available literature data (35), *K*_DI_ was assumed to be 70-fold less than *K*_Do_ in the model. Then, using the relation, *r_x_* = *k*_x_ * *K*_D**o**_, we optimized *k_i_d_* and *k_id_* for open-inactivated state of neutral and charged drugs (4).

### Computed dofetilide concentrations

We used the population *C*_max_ (maximum plasma concentration) of dofetilide: 2.72 ng/mL (33), and converted it to nanomolar (nM) concentration 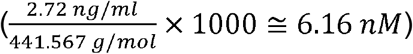 in the models, where 441.567 g/mol is dofetilide molar mass.

### Simulation of TRIaD in dofetilide and control case in O’Hara-Rudy Human model

First, action potential duration (APD) *Triangulation* was calculated as the repolarization time from APD_30_ to APD_90_ from 1000 simulated cells with application of noise current. The noise current was calculated using the equation from (36),

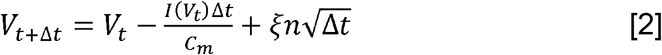

where *n* is a random number between 0 and 1 from a Gaussian distribution, and Δ*t* is the time step. *ξ* (= 0.3) is the diffusion coefficient, which defines the amplitude of noise (36). The noise current was generated and applied to the membrane potential *V_t_* throughout the simulated time course. *Reverse-use-dependence* was measured as APD_90_ at steady state for each pacing cycle length (from 2 Hz to 0.5 Hz), and APD adaptation curves were constructed. *Beat-to-beat (bTb) Instability* was simulated by applying small amplitude inward currents randomly between −0.1 to −0.2 pA/pF for 50 ms over the course of the action potential plateau between 10 to 700 ms at 1 Hz for 1000 beats.

### Fiber simulations

We simulated a transmural fiber composed of 165 O’Hara-Rudy human ventricular cells (37) (Δ*x* = Δ*y* = 100 μm) connected by resistances to simulate gap junctions (38). The fiber contains an endocardial region (cells 1 to 80) and epicardial region (cells 81 to 165), which showed a linear decrease in APDs (39, 40). *G*_Kr_ was monotonically increased from 0.04 to 0.05. The heart was paced at 1 Hz to match the clinically observed QT intervals ~400 ms (41–43). AP simulations were carried out in epi/endocardial cells by changing various ion channel conductances (37). The stimulus is applied to the first cell.

The fiber was paced at varying cycle length from 800 to 1400 ms for 200 beats (mean heart rates = 61 bpm) in order to match the clinical data (56.8±6.4 bpm) (33). Pseudo ECGs were computed from the transmembrane potential *V_m_* using the integral expression as in Gima and Rudy (44). Heart rate corrected QT (QTc) was computed using Fridericia formula using the cubic root of RR interval (45).

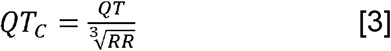

*Spatial APD dispersion* was measured using the T-wave area indicator, which was calculated as the T-wave amplitude on the computed pseudo-ECGs. For this purpose, a 1-dimensional model of the transmural wedge preparation, as described in (5), was stimulated by applying a standard short-long protocol as follows: The transmural wedge preparation was stimulated by a train of pulses (S1) at 1 Hz pacing rate until the steady-state was reached followed by a premature beat (S1-S2 interval = 800 ms) and then a delayed beat (S3) was delivered after a long pause (S2-S3 interval = 5000 ms). T-wave area calculations were computed as follows:

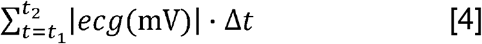

where Δt =1 ms, *t*_1_ is the time where ECG equals to T_peak_ – 0.9*(T_peak_ = minimum of left side of T wave), and *t*_2_ is the time where ECG equals to T_peak_ – 0.9*(T_peak_ = minimum of right side of T wave).

### Frequency-dependent QT prolongation

The fiber was paced at 1Hz for 1000 beats (S1) and then a second stimulus (S2) was applied after a varying RR interval (between 550 and 1200 ms). The QT interval, in response to S2, was recorded. The same simulations were carried out 6 times for both control and 2.72 ng/mL dofetilide with noise currents, and the relative changes in slope of relationship of QT and preceding RR intervals were calculated.

### Two-dimensional simulations

2D simulations were performed to determine if proarrhythmic phenomena observed in lower dimensions cause reentrant arrhythmias. Current flow is described by the following equation:

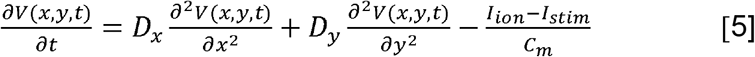

where *V* is the membrane potential, *x* and *y* are distances in the longitudinal and transverse directions, respectively, *D_x_* and *D_y_* are diffusion coefficients in the *x* and *y* directions. We simulated a *heterogeneous* and a *homogenous* cardiac tissue on a 500 by 500 pixel grid with Δ*x* = Δ*y* = 100 μm. The heterogeneous tissue contains an endocardial region (fibers 1 to 180) and epicardial region (fibers 181 to 500). We also incorporated anisotropic effects by setting *D_x_* and *D_y_* such that the ratio of conduction velocities is 1:2 (46). A typical S1-S2 protocol was used for Figure 5. The tissue was first paced (S1) in a 0.5 cm × 1.1 cm area on the left edge of the endocardial region, and a premature stimulus (S2) was then applied in a 1.8 cm × 1.5 cm area on the top left corner of the endocardial region. Small amplitude inward currents were randomly applied between −0.1 to −0.45 pA/pF on *each cell* in both *heterogeneous* and *homogenous* tissues after 0.5 ms.

### Action potential duration (APD) mapping

We reconstructed the “human transmural myocardial wedge” based on data describing transmural action potential heterogeneity mapped from normal human left ventricle (40) (Figure 6A). First, the O’Hara-Rudy human model (37) was used to generate a *G*_Kr_ lookup table corresponding to APD_80_. Next, experimental 2D APD_80_ map (100 × 100 – Figure 6A) was used to create a 2D *G*_Kr_ map using the *G*_Kr_ lookup table. Then the twodimensional *G*_kr_ values (100×100) were used to simulate APD_80_ at pacing rate of 0.5 Hz. We then constructed ***3D wedge*** of 100 by 100 by 1 cells with Δ*x* = Δ*y* = 200 μm and Δ*z* = 500 μm using this APD mapping data. Current flow is described by the following equation:

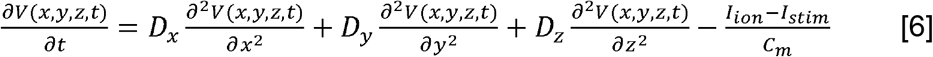

Where *V* is the membrane potential. *D_x_, D_y_* and *D_z_* are diffusion coefficients in the *x, y* and *z* directions. Stimulus current *I_stim_* is 150 mA/cm^2^ for 0.5 ms. We also incorporated anisotropic effects by setting *D_x_, D_y_* and *D_z_* such that the ratio of conduction velocities is 2:4:1 (46).

### Local sensitivity analysis

We calculated elasticity coefficients (sensitivities) for arrhythmia vulnerability parameters from the TRIaD based simulations. The protocol for arrhythmia vulnerability parameters from the TRIaD based simulations is the same as in Figure 5 in the presence of 2.72 ng/mL. The relative change in each arrhythmia vulnerability parameter in response to a parameter perturbation (local sensitivity) was calculated by following equation:

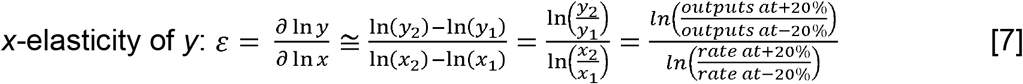

In each case, the model rate constants were increased and decreased by 20%. If |*ε*| > 1 (sensitive), *y* (model output) changes more than changes in *x* (the model rate). In contrast, |*ε*| < 1 (insensitive) indicates that *y* (model output) changes less than changes in *x* (the model rate) (47, 48).

### Simulated data of the arrhythmia vulnerability parameters from TRIaD for random forest machine learning application

*1) APD TRI:* APD triangulations were calculated from APD_30_ to APD_90_ from 1000 simulated cells with noise currents. The protocol is same as in **Simulation of TRIaD**. *2) bTb instability:* 1000 simulated APDs_90_ were recorded by adding noise currents into membrane potential calculations. The protocol is same as in **Simulation of TRIaD**. *3) RUD:* APDs_90_ were recorded from 1000 cells at slow pacing rate (BCL = 2000 ms) with noise currents. *4) T-wave area:* The transmural fibers were stimulated by a standard short-long protocol, and T-wave areas were calculated (as described in **Fiber simulations**) from 1000 cases with noise currents.

### Random forest machine learning algorithm

We applied a multivariate correlation-based filter selection (CFS) technique using a random forest machine learning algorithm for multiclass classification (49) to indicate the importance of each arrhythmia vulnerability parameter from **TRIaD** (*APD TRI, bTb instability, T-wave area and RUD*) towards the target (classification of control and different dofetilide doses). We calculated the existing correlation coefficients using Pearson’s correlation coefficient Eq. [8] to explore the linear dependence of arrhythmia vulnerability parameters from **TRIaD**. Pearson’s coefficient values vary between −1 and 1, where 1 is highly correlated (changes in *x*_1_ are correlated with changes in *x*_2_) and −1 is highly anticorrelated (changes in *x*_1_ are negatively correlated with changes in *x*_2_) (50).

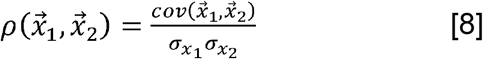

where 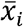 is a vector consisting of each **TRIaD** parameter’s observations, *cov* is covariance between two **TRIaD** parameters and *σ* is standard deviation of each **TRIaD** parameter observed in the simulations.

Random forest is an ensemble decision tree, which contains a collection of single decision trees. Each tree is built over random extraction of the observations from the dataset and the random extraction of the features (arrhythmia vulnerability parameters in this case) (51). For each feature a series of questions are formulated so that the answer to those questions lead to the best possible separation of classes into groups that contain only one class or the majority of one class at each node. Therefore, the importance of each feature is indicated by purity in each branched out node, and impurity within a node for a particular feature indicates reduced importance. The importance of each feature is averaged across all the trees in random forest classifier to determine the final importance of the features (52).

In order to train the random forest classifier, we first performed feature scaling using Eq. [9] to assure that each feature in the dataset has zero-mean and unit-variance (53)

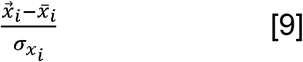

After feature scaling, we split our dataset into random training and testing subsets (using 80% of data for training and 20% of data for testing the performance of the trained classifier). Then we trained the random forest multiclass classifier using the training subset and set 10 trees and entropy at the same time as the measure of impurity for the random forest classifier. Entropy is a measure that controls how the decision tree decides which feature to choose first and where to split the data (Eq. [10]).

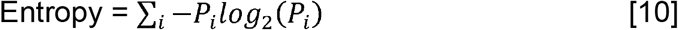

Where the *P_i_* is the fraction of examples in class *i*.

## RESULTS

The first step we took to build a computational pipeline for predictive cardiac safety pharmacology was to compute numerical estimates of drug-binding energetics and kinetics from atomic scale simulations of drug interactions with our hERG channel model in an open state (Figure 1A – C).

One-dimensional PMF (free energy) and diffusion coefficient profiles (shown in Fig. 1B), computed from umbrella sampling MD simulations, were used to calculate the binding free energies and diffusion rates of dofetilide into the open state of the hERG pore. These values allowed for the constrained optimization of rate constants that were used to populate a state dependent hERG function scale model shown schematically in Figure 1D. To simulate drug interactions with hERG, we used drug diffusion rates D (~0.11×10-6 cm2/s and ~0.15×10-6 cm2/s for neutral and charged) and affinities (dissociation constants *K*_Do_, Figure 1C) from the PMF calculation to constrain the drug “on” ([drug]\**k_x_*) and “off” (*r_x_*) rates for neutral and charged drugs. The details of the calculations are presented in (REF). The “off” rate from the open state was calculated as *r_x_* = *k_x_* * *K*_Do_, where *K*_Do_ was computed using PMF profiles from the atomistic scale MD simulation (Figure 1C). The “on” rates for open-inactivated state were optimized to the experimentally obtained IC_50_ curve from Vicente et al. (33) by assuming that dofetilide binds 70-fold stronger to this channel state, i.e. *K*_DI_ = (*K*_Do_ / 70) (35), as shown in Figure 1D (bottom). In both experiment and simulations (Figure 1E), peak *I*_Kr_ was recorded at the end of the 3-s activating step to 0 mV with drug concentrations from 0 to 7.5 ng/mL. Percentage of drug block was calculated by (*I*_control_ – *I*_drug_)*100/*I*_control_ and compared to experimental data (33), demonstrating good fit (Figure 1E).

Next, we subjected our computational pipeline for safety pharmacology to a validation test using the gold standard data: human clinical data in the form of electrocardiograms in the absence and presence of dofetilide. To do so, as shown schematically in Figure 2A, we incorporated the function scale model of dofetilide interaction with the hERG channel (Figure 1D) into the O’Hara-Rudy human cardiac ventricular myocyte model and then extended this model to construct a one-dimensional strand of O’Hara-Rudy cells (37) by connecting them via simulated resistive elements to represent gap junctions. We applied a simulated stimulus current at one end to initiate a propagating one-dimensional wave at the clinically reported heart rate between 43 and 75 beat per minute (bpm) (33). Figure 2B shows the calculation of the spatial and temporal gradients of electrical activity used to construct a heart rate corrected pseudo ECG (QT_C_ interval) for a range of dofetilide concentrations.

**Fig. 2.**
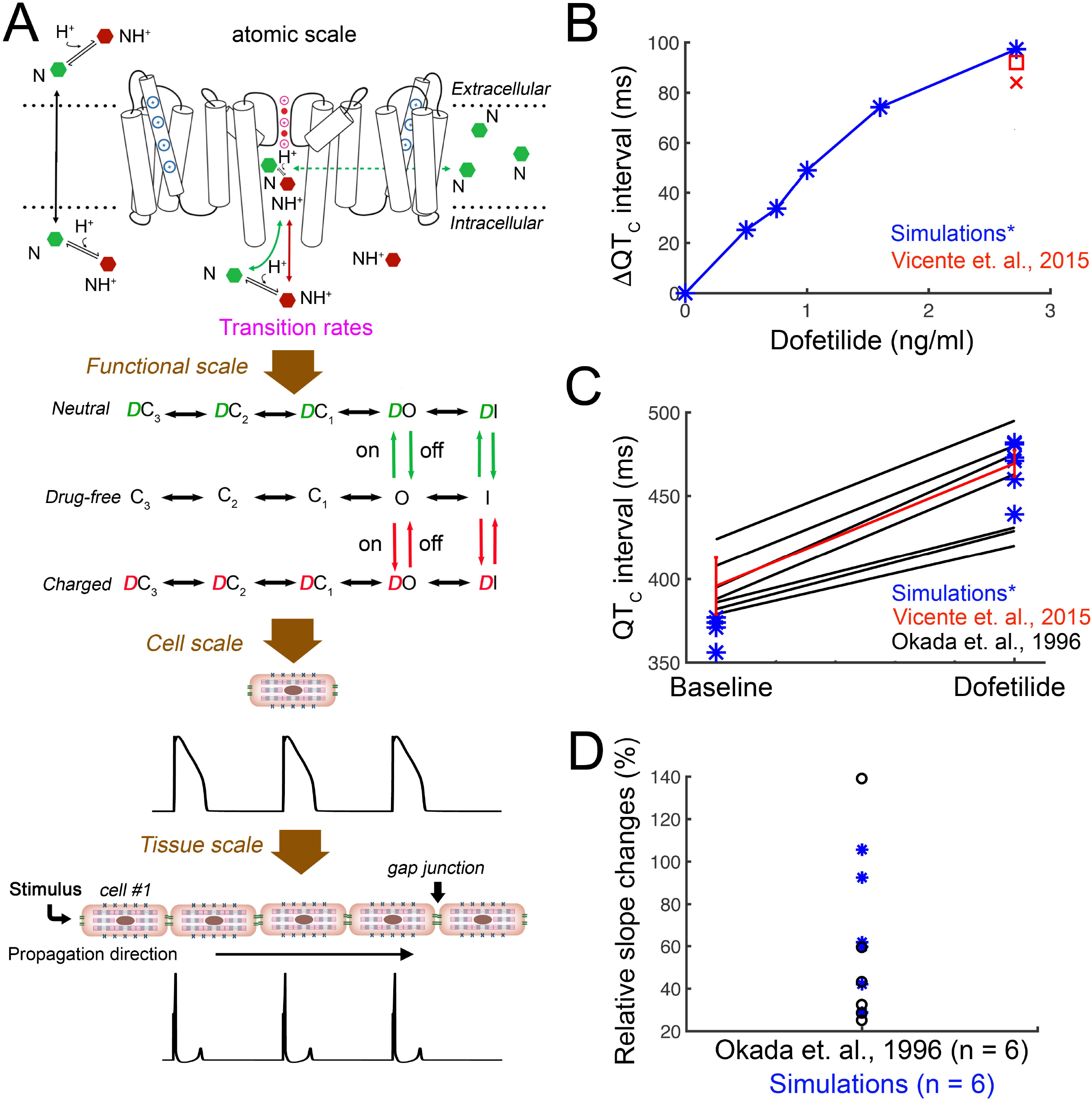
Validation of the dofetilide computational screening pipeline prototype with human clinical data. A) Schematic diagram presents the connection of atomistic scale model of dofetilide interaction with the hERG channel (top) to the corresponding functional protein scale model (middle) used in single-cell to tissue level simulations and computed pseudo ECG (bottom). B) A heart rate corrected pseudo ECG (ΔQT_C_ interval) was computed from a 1-dimensional strand of O’Hara-Rudy human cardiac ventricular myocytes for pacing frequencies between 43 – 75 bpm for a range of dofetilide concentrations (blue) compared to clinical data (red). (C) Comparison of human clinical data showing control and dofetilide affected rate corrected QT intervals (9, 33) (black and red lines) and simulated mean values under the same conditions (blue asterisks). Red line: two subjects received a single dose of 0.5 mg (population’s mean maximum concentration C_max_ is 2.7±0.3 ng/mL). Blue asterisks (*): concentration 2.72 ng/mL (~ 6.16 nM) was used in the simulations. Black lines: subjects received 0.5 to 0.75 mg twice a day. (D) The clinically observed and *in silico* prediction of QT intervals over a wide range of preceding RR intervals after 2.72 ng/mL dofetilide application. Rate dependent changes in the QT interval were tracked as the slope of the linear regression line estimating the 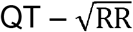 relation.

Figure 2C shows the comparison of human clinical data under drug free conditions and following application of 2.72 ng/mL dofetilide (9, 33). The simulated mean values compared to clinically obtained data from humans are in excellent agreement, thereby providing an indication of the validity and predictive value of the computational pipeline to recapitulate the effect of a drug on the human QT interval.

Finally, as shown in Figure 2D, we simulated QT intervals over a wide range of preceding RR intervals after 2.72 ng/mL dofetilide application and compared to the clinically observed changes (9). Each cell in the simulated tissue was subjected to a physiological noise current in order to introduce physiologically relevant variability. Rate dependent changes in the QT interval were tracked as the slope of the linear regression line estimating the 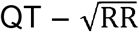 relation. Again, the predicted relationship falls within the range of clinical data, indicating that the model can reproduce rate dependent changes in drug-induced QT interval.

We next carried out computational screening in O’Hara-Rudy human computational ventricular myocytes for the effect of dofetilide to promote proarrhythmia by tracking the parameters comprising the **TRIaD**. We tracked each parameter in the absence of drug (control conditions in black) and in the presence of average patient plasma concentration of 2.72 ng/mL dofetilide (red). This approach was also carried out in simulated rabbit ventricular myocytes to explore the effects of species differences. The predicted results in rabbit were very similar to human and are shown in Supplemental Figure S1.

In Figure 3A, ***temporal APD dispersion*** was quantified in a cell population of 1000 individual simulated cardiac myocyte action potentials constructed by incorporating physiological noise (36, 54, 55). APD temporal dispersion was quantified as the difference between the maximum and minimum APD. Dofetilide within the clinical dosing range has a clear effect to promote temporal APD variability in the presence of the drug (Control – 47 ms; 2.72 ng/mL Dofetilide −78 ms). Figure 3B illustrates the effect of dofetilide to promote ***triangulation*** of the action potential as a function of APD prolongation. In the absence of drug, control cells had a slope of 0.25, while 2.72 ng/mL Dofetilide increased the slope to 0.66. Figure 3C shows Poincaré plots of sequential APD pairs indicating beat-to-beat ***instability*** following the application of small electrical perturbations in the absence of drug or with 2.72 ng/ml dofetilide. Instability was assessed by applying small amplitude inward currents randomly between −0.1 to −0.2 pA/pF for 50 ms over the course of the action potential plateau at a basic cycle length of 1000 ms. In Figure 3D, ***reverse use dependence*** induced by dofetilide was evaluated. The action potential adaptation curves were generated using APD_90_ values from human computational ventricular myocytes at steady state at the indicated pacing frequencies. When dofetilide (red) was applied, there was a clear steepening of the APD adaptation curve compared to the baseline drug-free case (black). Panel E shows spatial dispersion of APD that was quantified in tissue by integrating the area under predicted T-wave following a long pause (5000 ms). See methods section for details. The table in Panel F shows the quantified increase in area (79%) under the T-wave when dofetilide is applied.

**Fig. 3.**
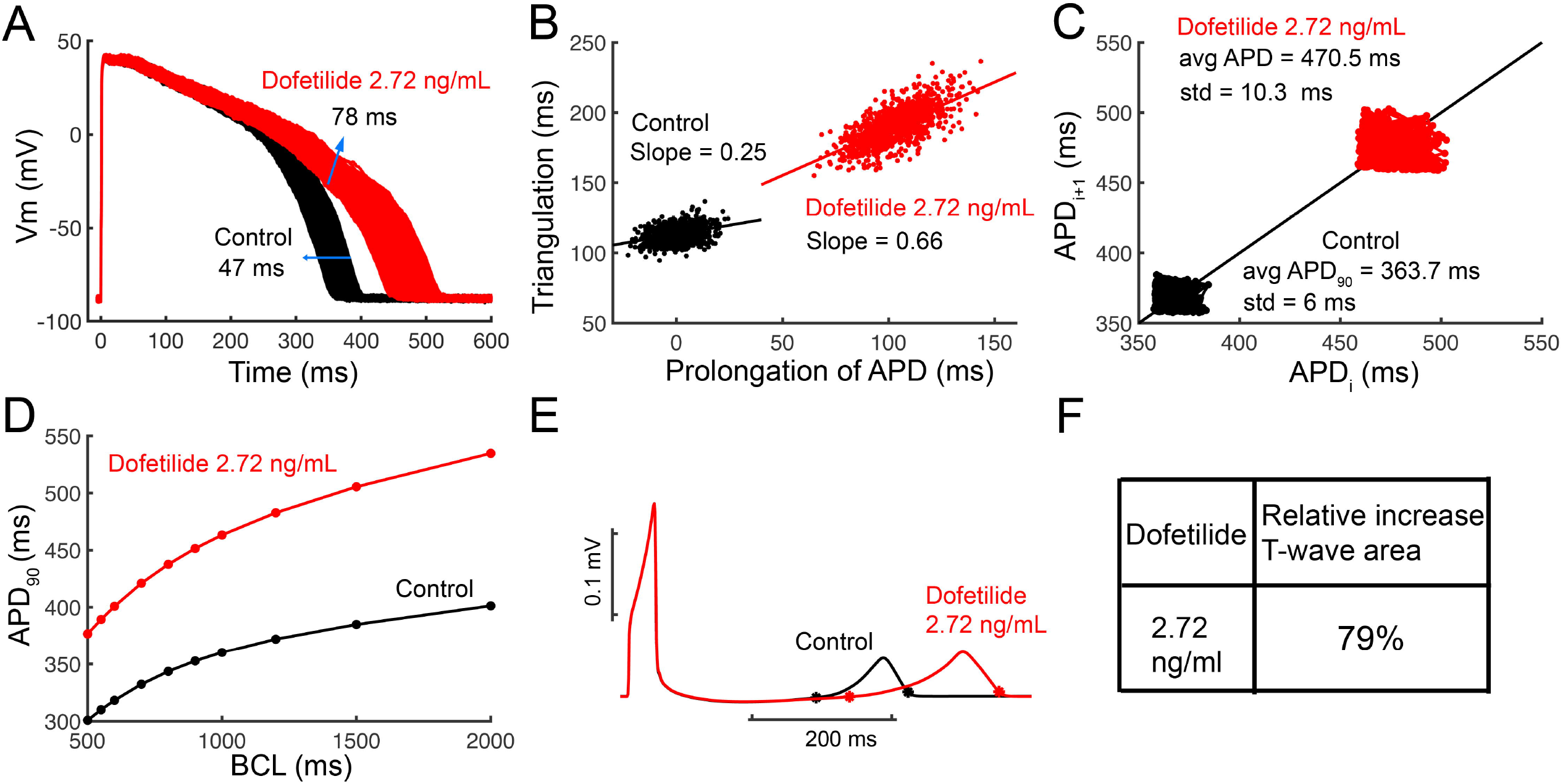
Computational screening for dofetilide induced arrhythmia vulnerability. A) Predicted *temporal APD dispersion* of 1000 simulated O’Hara-Rudy human ventricular action potentials generated after incorporating physiological noise to induce beat-to-beat variability at 1 Hz in the drug-free control case and following simulated application of Dofetilide (2.72 ng/mL). *Dispersion* of APD was quantified as the difference between the maximum and minimum of 1000 individual cells (Control – 47 ms; Dofetilide 2.72 ng/mL = 78 ms). B) Action potential *triangulation* as a function of APD prolongation for individual cells for control (slope = 0.25), and Dofetilide 2.72 ng/mL (slope = 0.66) conditions C) Simulated beat-to-beat *instability* of action potentials to small perturbations before and after application of drugs. Poincaré plots of sequential APD pairs indicating beat-to-beat instability are shown. D) Action potential adaptation curves show APD_90_ at various pacing frequencies with or without dofetilide, demonstrating drug reverse use dependence. E) pseudo ECGs after a long pause (5000 ms) are shown for control and dofetilide 2.72 ng/mL conditions. F) Relative increased T-wave area by 79% with dofetilide 2.72 ng/mL.

Arrhythmia is fundamentally an emergent spatial phenomenon. Accordingly, simulations were performed via an *in silico* diagnostic protocol by applying spatial noise to promote electrical instability and determine if dofetilide promotes arrhythmia triggers and/or proarrhythmic rhythms under these vulnerable circumstances. Two-dimensional homogeneous (Panels A and B, endocardial cells) and heterogeneous (Panels C and D, endocardial region (cells 1 to 180) and epicardial region (cells 181 to 500)) anisotropic human ventricular *in silico* tissues (5 cm × 5 cm) with a linear decrease in APD as indicated by experimental data (39, 40) were simulated. Each simulated tissue contained randomized spatial heterogeneity imposed by the application of low amplitude perturbations in the form of small inward currents, which were randomly applied between −0.1 to −0.45 pA/pF to ***each cell*** in the tissue at each time step for the duration of the simulation. The results are shown in Figure 4. In Panels A and B, the homogeneous tissue simulations showed that in the absence of drug, the tissue was very electrically stable, and normal cellular and tissue behavior is observed. However, the presence of 2.72 ng/ml dofetilide resulted in the emergence of early afterdepolarizations (EADs) in some cells and not others, resulting in spatial dispersion of repolarization. As shown in Panels C and D, the effect persisted when the tissue was heterogeneous (to mimic transmural heterogeneity), with considerably reduced dispersion of repolarization (as epicardial cells fire last, but repolarize first), and spatial repolarization gradients were observed in the setting of dofetilide only. These simulations suggest that in the presence of low levels of electrical instability as are likely to occur in a variety of disease settings, the application of a clinically relevant dose of dofetilide promotes clear spatial dispersion of repolarization.

**Fig. 4.**
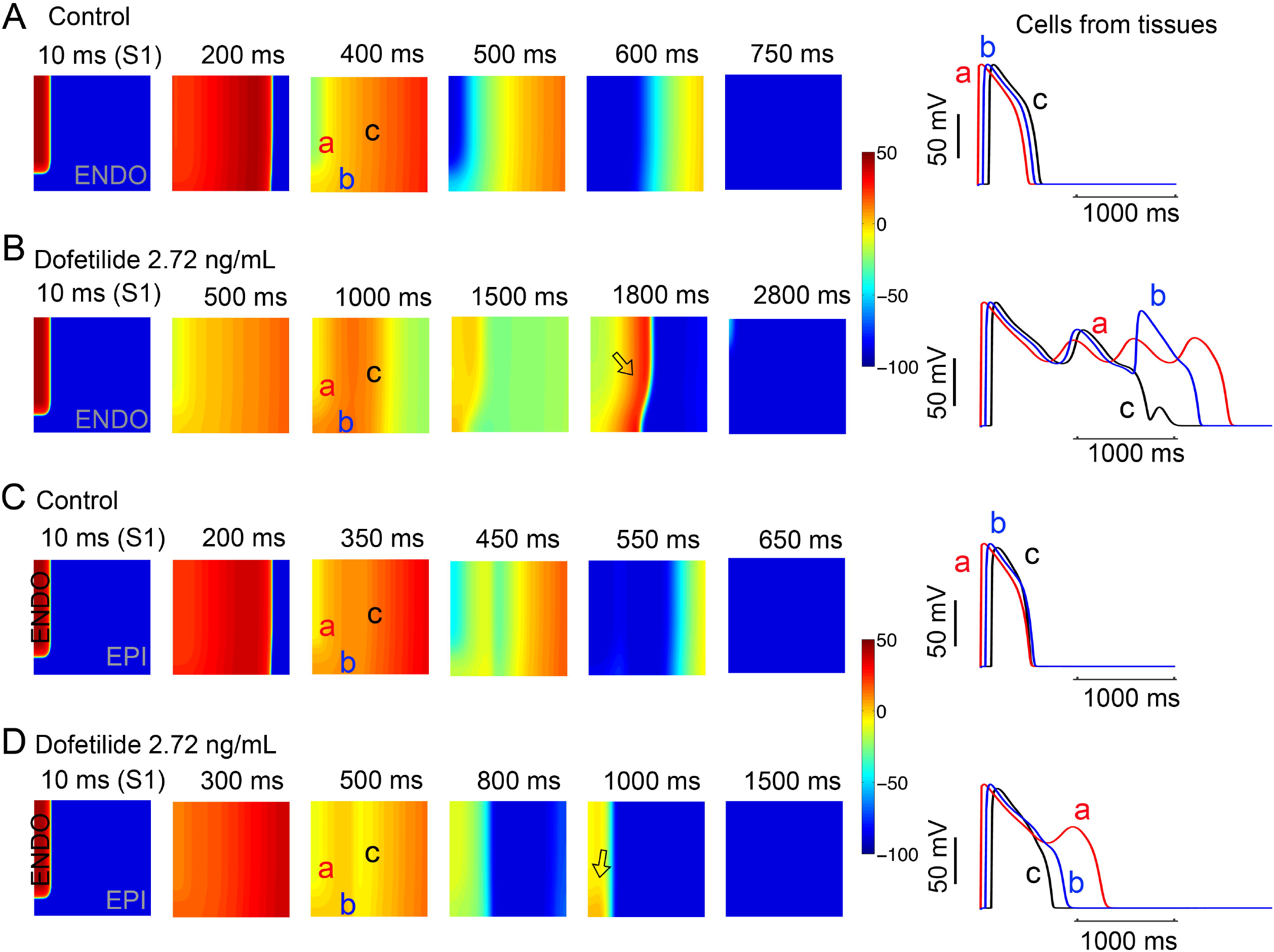
*In silico* diagnostic test in tissue reveals arrhythmia triggers with dofetilide. Time snapshots (colored boxes) with voltage gradients are shown for twodimensional simulated tissue as described in methods. Membrane voltages are indicated by the color gradient. Two-dimensional homogeneous (Panels A and B, endocardial cells) and heterogeneous (Panels C and D, endocardial region (cells 1 to 180) and epicardial region (cells 181 to 500)) anisotropic human ventricular *in silico* tissue composed of (5 cm × 5 cm) simulated myocytes. Single APs from sites ‘a’, ‘b’, and ‘c’ in the simulated tissues are shown in the right panels for each case.

We next set out to test the effect of dofetilide in the setting of extrasystolic excitable triggers in the heterogeneous tissue as shown in Figure 5. Again, in the presence of spatial noise to promote low-level electrical instability, when 2D tissue was simulated using a typical S1-S2 protocol (56, 57), 2.72 ng/ml dofetilide application resulted in considerable dispersion of repolarization that was absent in control drug free tissue. The tissue was first paced (S1) (first panel) in a 0.5 cm × 1.1 cm area on the left edge of the endocardial region, and a premature stimulus S2 (third panel) was then applied in a 1.8 cm × 1.5 cm area on the top left corner of the endocardial region. As described above, spatial heterogeneity was applied via small amplitude inward currents randomly applied between −0.1 to −0.45 pA/pF to each cell in heterogeneous tissues after 0.5 ms. Time snapshots (panels) with voltage gradients indicated by the color map are shown in Fig. 5. These maps were constructed following the last planar wave (S1) (first panel) and throughout termination of the most persistent wave after S2 stimulus (last panel). The corresponding action potentials from three points in space are shown in the right panels. In the absence of drug (top row), there was no persistence of electrical instability. In the bottom row the effect of dofetilide is shown, which reproducibly (*n* = 5 simulations) promoted numerous persistent arrhythmia triggers observed as afterdepolarizations in the cellular action potentials (right).

**Fig. 5.**
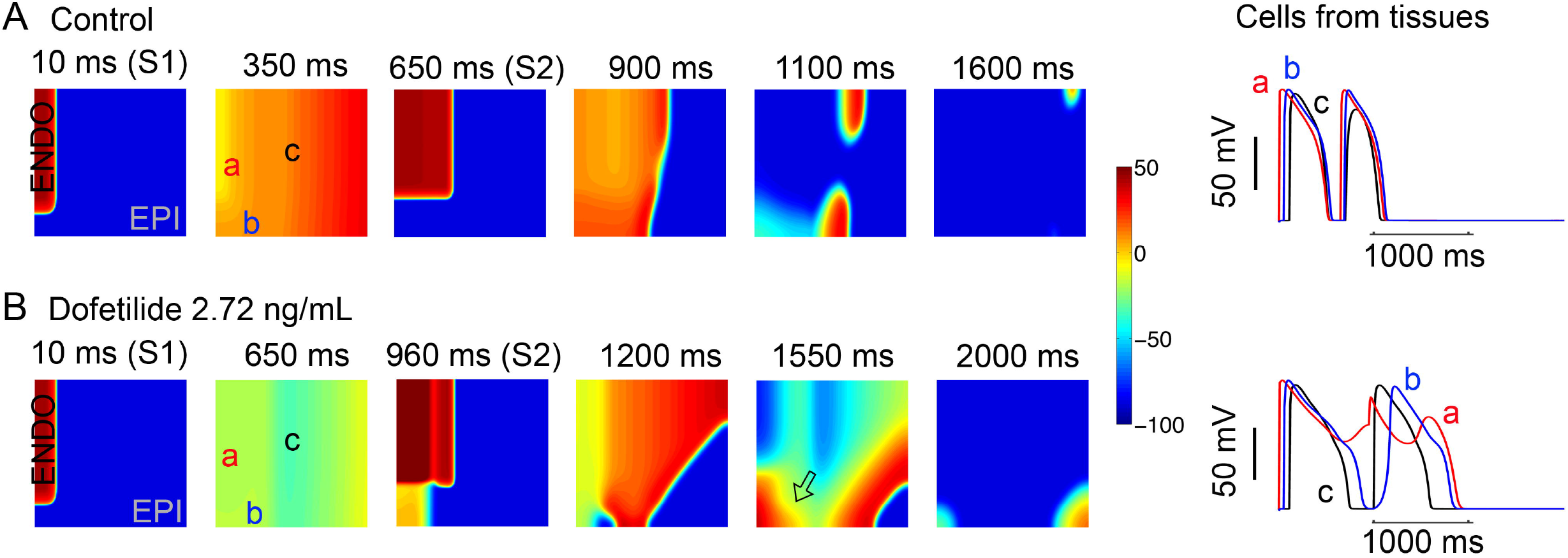
*In silico* diagnostic test to reveal vulnerability to torsades de points arrhythmias by extrasystoles. Time snapshots (colored boxes) with voltage gradients are shown for two-dimensional simulated tissue as described below. Membrane voltages are indicated by the color gradient. The corresponding action potentials from three points in space (‘a’, ‘b’, and ‘c’) are shown in the right panels. A) In the absence of drug (top row), arrhythmia triggers were not inducible. B) In the bottom row the effect of dofetilide is shown: numerous arrhythmia triggers observed as afterdepolarizations and spatial repolarization gradients in underlying cellular action potentials (right). Single APs from site ‘a’, ‘b’, and ‘c’ in the simulated tissues are shown in the right panels for each case.

We eveloped an *in silico* left ventricular (LV) 3D wedge reconstruction of the human cardiac tissue based on the experimental data from Glukhov et al. (40) for the drug free case and with 2.72 ng/ml dofetilide applied. Clinical and experimental data suggest that male and female subjects respond differently to dofetilide intervention, with females exhibiting increased sensitivity and proclivity to abnormal rhythm (58–60). The LV wedges were paced at basic cycle of 1000 ms for 5 beats with small amplitude inward currents randomly applied between −0.1 to −0.45 pA/pF to *each cell* after 0.5 ms. The model simulations show drug-induced QT interval prolongation both in male and female. However, in females, dofetilide caused considerably larger prolongation of the QT interval, at 60 bpm, as well as at 80 bpm.

In order to assess the specific drug – channel interactions that comprise the dofetilide structure-activity relationship and the link to proarrhythmia, we undertook sensitivity analysis to determine how sensitive the arrhythmia vulnerability parameters from the TRIaD based simulations are to our underlying model parameters (Figure 7). As shown in Figure 7A, we first carried out an *in silico* test of the local sensitivity of the slope of the relationship between action potential *triangulation* and APD prolongation in O’Hara-Rudy computational myocytes plotted for a range of drug “on” (*k_o_d_, k_od_, k_i_d_*, and *k_id_*) and “off” (*r_o_d_, r_od_, r_i_d_* and *r_id_*) model transition rates for open and open-inactivated states by increasing and decreasing each rate at ±20% for neutral (green) and charged (red), and calculated elasticity coefficients. The local sensitivity analysis showed that perturbation to the rate constant *k_i_d_* (open-inactivated state binding) of neutral drug results in the greatest effect on the slope of APD triangulation. The elasticity value of *k_i_d_* shown in Figure 7A is 0.52, which suggests that if the rate constant *k_i_d_* is increased by 1% then the slope of APD triangulation increases by approximately 0.52%, indicating relative insensitivity to even the most sensitive model parameter. Perturbation to all other rate constants also results in less than 1% changes in the slope of APD triangulation. Similarly, in Figure 7B, simulated beat-to-beat *instability* of action potentials (average and standard deviation of APD_90_ for each rate perturbation are shown) with respect to transition rate changes are demonstrated. And Figure 7C shows relative changes in transition rates with respect to the APD_90_ at a slow pacing rate, which reflect maximal reverse use dependence (BCL = 2000 ms). Again, binding and unbinding rate parameters of neutral drug (green bars) for inactivated channel state caused more changes in the model outputs compared to other parameters, but the model outputs are still relatively insensitive. Finally, Figure 7D, shows higher elasticity values of the T-wave area to neutral drug (green bars) binding and unbinding rates for inactivated state. However, the analysis showed that the model was robust to perturbations (elasticity value < 1).

**Fig. 6.**
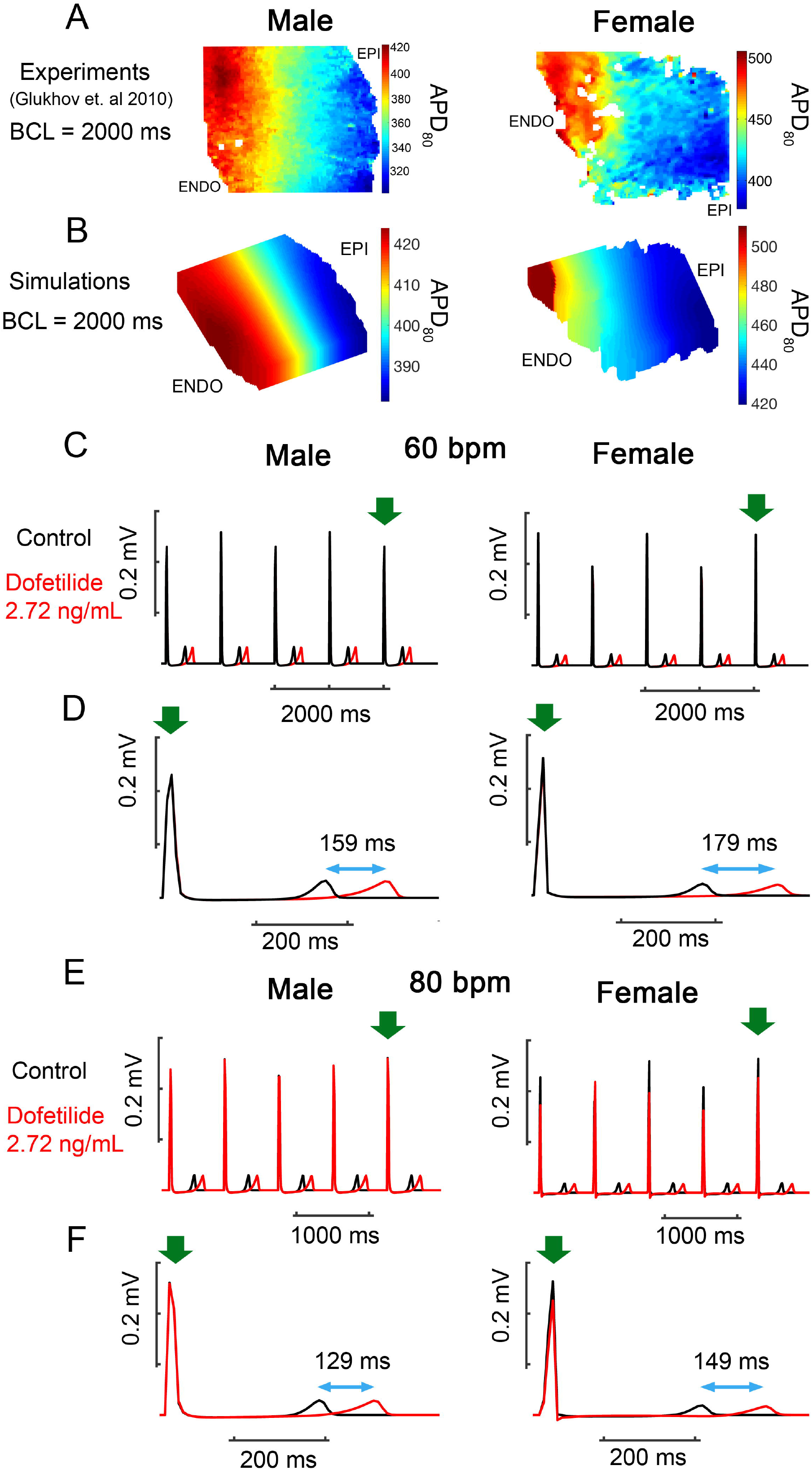
In silico 3D reconstructed left ventricular (LV) wedge based on Glukhov et al. (40) paced with 2.72 ng/mL dofetilide applied. Experimental data from normal human female (right) and male (left) left ventricle (40) shown in panel A. (B) Reconstructed a human tissue in silico from these data from normal explanted heart at a pacing cycle length of 2000 ms (male – left and female – right). Pseudo ECGs show the effect of dofetilide applications on QT intervals for male (left) and female (right). (C – D) The LV wedges were paced at 60 bpm (basic cycle of 1000 ms) for 20 beats with noise applied (small amplitude inward currents randomly applied between −0.1 to −0.45 pA/pF on *each cell* after 0.5 ms). The last 5 beats were shown on the top panel (male – left and female – right). Green arrow is the final stimulus. (E – F) The wedges were paced at 80 beat per min (basic cycle of 750 ms) for 20 beats with noise applied (small amplitude inward currents randomly applied between −0.1 to −0.45 pA/pF on *each cell* after 0.5 ms). The last 5 beats were shown on the top panel (male – left and female – right). Green arrow is the final stimulus.

**Fig. 7.**
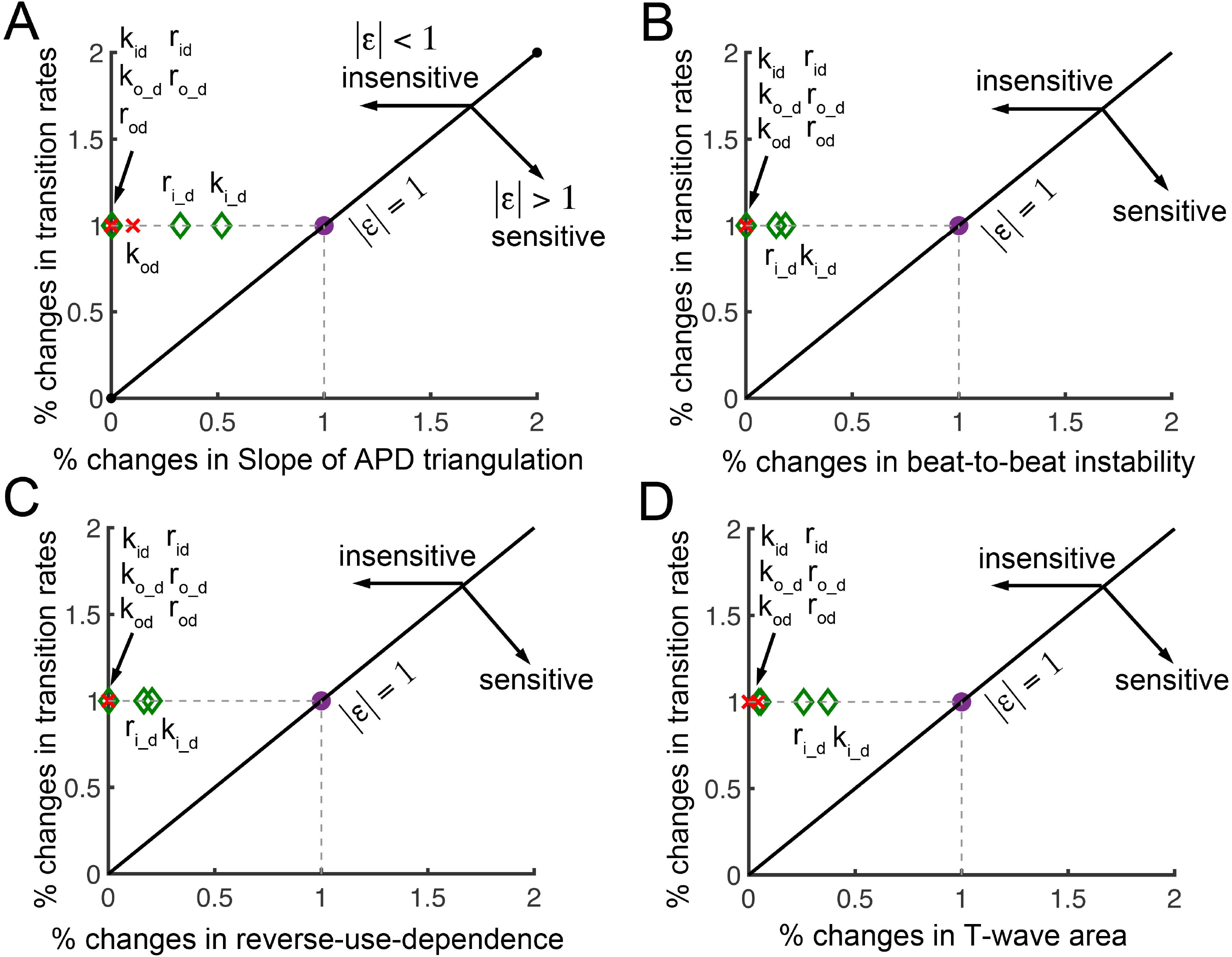
Local sensitivity analysis of arrhythmia vulnerability parameters from the TRIaD. A) The sensitivity of the slope of the relationship between action potential *triangulation* in O’Hara-Rudy computational myocytes plotted for a range of drug “on” (*k_o_d_, k_od_, k_i_d_*, and *k_id_*) and “off” (*r_o_d_, r_od_, r_i_d_* and *r_id_*) model transition rates for open and inactivated states by increasing and decreasing each rate at ±20% for neutral (green) and charged(red). B) Sensitivity of simulated beat-to-beat *instability* of action potentials for a range of rate constants. C) The sensitivity to changes in charged and neutral drug transition rates of the recorded APD_90_ at BCL of 2000 ms. D) Sensitivity of the T-wave area to model transition rates.

Finally, we used random forest machine learning algorithm to evaluate the importance of each arrhythmia vulnerability parameter from the **TRIaD** (*APD TRI, bTb instability, T-wave area and RUD*) to determine the final target classification (control versus drug affected at different dofetilide doses). In general, to make a good classification algorithm it is important to identify parameters that are correlated with the target output of interest but uncorrelated among themselves. Therefore, we used Pearson’s correlation coefficient (see methods) to explore the linear dependence of **TRIaD** parameters in Figure 8A. If two **TRIaD** parameters are highly correlated, they provide redundant information. As shown in Figure 8A, all **TRIaD** parameters are highly correlated. After finding highly correlated features, we constructed a multiclass random forest classification (see methods) using all the correlated **TRIaD** parameters. The random forest classifier outputs the importance of all **TRIaD** parameters and indicates the importance of each parameter to correctly predict the target (drug free, or low, medium or high risk (indicated by dose)). Figure 8B suggested that beat-to-beat (bTb) *instability* is the most important feature that contributes to classify control case versus drug affected case. Figure 8C illustrates the multiclass classified region for control, dofetilide 0. 1 ng/mL, dofetilide 1 ng/mL and dofetilide 2.72 ng/mL using *bTb instability* and *RUD* with random forest classification’s accuracy of 93%. Please have Parya respond to Igor’s comment

**Fig. 8.**
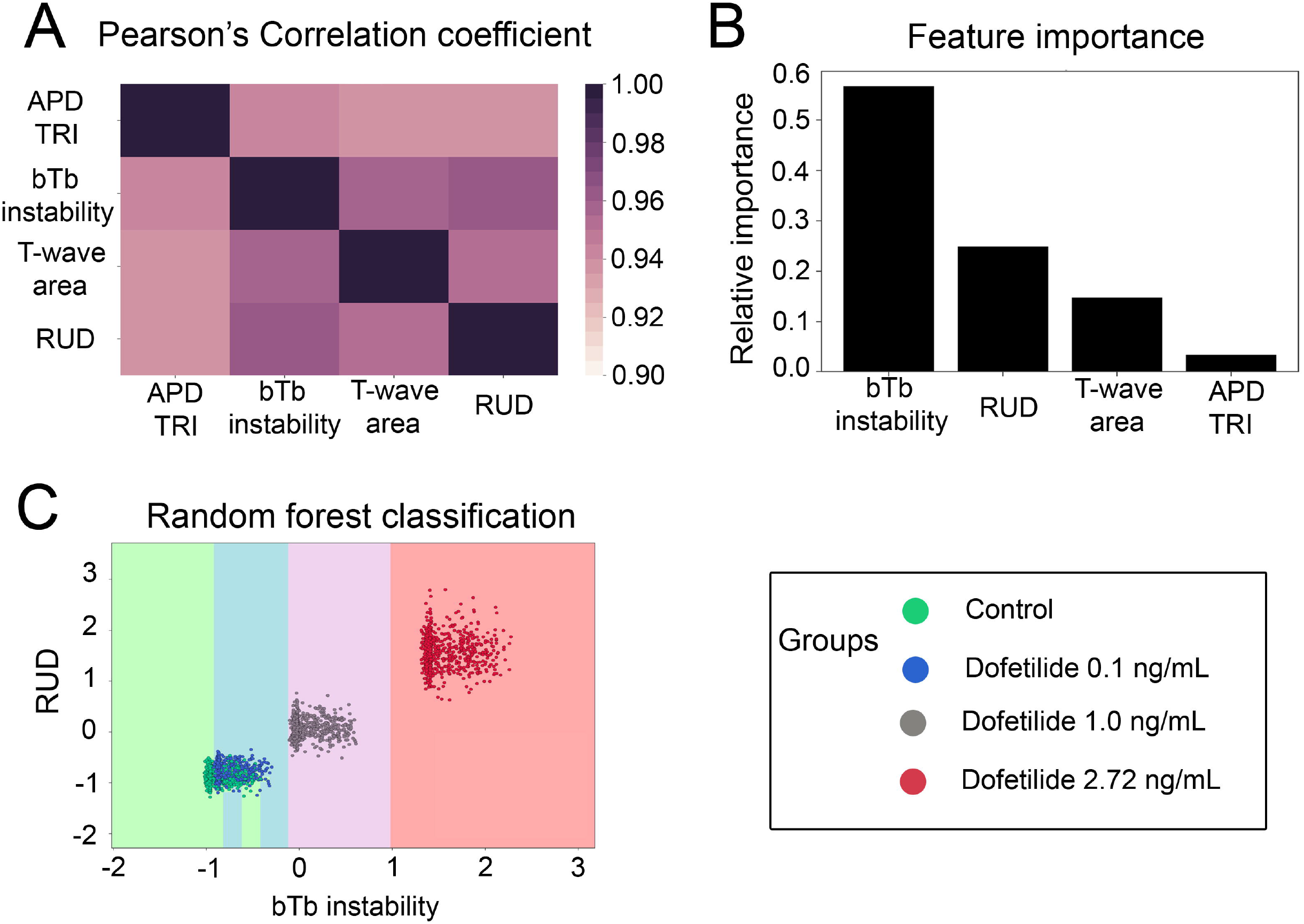
Application of random forest machine learning to classify dose-dependent importance of TRIaD-linked parameters to arrhythmia emergence. A) The Pearson’s correlation coefficients between paired TRIaD parameters. Pearson’s coefficient value indicated by the color gradient, where 1 is highly correlated (black). B) Beat-to-beat (bTb) *instability* emerged as the most important feature from the TRIaD parameters. C) Classified regions for control, dofetilide 0.1 ng/mL, dofetilide 1 ng/mL and dofetilide 2.72 ng/mL using beat-to-beat (bTb) *instability* and reverse use dependence (RUD).

## DISCUSSION

A major factor plaguing drug development is that there is no preclinical drug-screening tool that can accurately predict unintended drug induced cardiac arrhythmias from chemically similar drugs. The current approaches rely on substitute markers such as action potential duration or QT interval prolongation on the ECG. Unfortunately, QT prolongation is too sensitive and not specific in its indication of likelihood of arrhythmias. There is an urgent need to identify a new approach that can predict actual proarrhythmia from the drug chemistry rather than relying on surrogate indicators.

In this study we take the first steps to construct a computational pipeline for predictive safety pharmacology. The goal of this study was to develop a framework that will allow the detection of unsafe hERG blockers from their drug chemistry early in the preclinical screening process. Thus, we have assembled the process and utilized clinical data to demonstrate the utility for a proof-of-concept multiscale computational model to predict cardiac effects of dofetilide, a potent hERG blocker with a high pro-arrhythmia risk.

We began by developing physics based computer models (REF) that can account for channel conformation state and drug ionization state specific atomic-scale determinants of dofetilide interaction with hERG. This was accomplished through: 1) the development of open-state hERG atomistic structural model based on a recently published cryo-EM structure and its validation via all-atom MD simulations of K^+^ conduction through the channel pore under the applied voltage; 2) the development of empirical force field models for both charged and neutral forms of dofetilide and their validation via atomistic membrane partitioning MD simulations; 3) careful determination of free energy and diffusion coefficient profiles for drug binding to the channel pore using enhanced sampling all-atom MD simulations. See (REF) for more details.

### Molecular dynamics simulations yield novel insights of dofetilide interaction with hERG

Next, we utilized molecular dynamics (MD) simulations to predict association rates and affinities of dofetilide to the open state of the hERG K^+^ channel. Interestingly, this approach yielded novel information about the nature of dofetilide interactions with hERG. The molecular dynamics simulations suggest that neutral dofetilide preferentially interacts with the open channel state compared to the charged drug (see Figure 1B and 1C). Our previous function scale dofetilide model (32) was based on interpretation of experimental data, some of which suggested a 70-fold preferential binding to the inactivated state relative to the open state (35) (34) (61) (62). These data were used to estimate drug binding to the inactivated channel state in our model by assuming a 70-fold increase in predicted affinities from molecular dynamics simulations of the open state.

Interestingly, our previously published models required higher doses of dofetilide to cause prolongation of the QT interval (32), but the model generated from the molecular dynamics generated parameters in this study was able to reproduce dose-dependent prolongation of the QT interval in very close agreement to the clinical data (Figure 2B). These results serve as an important reminder for the difficulty in empirically deconstructing state-dependent drug mechanisms from experimental data. This is because the drug interaction often occurs on the same timescale (ms) as channel gating. In the physics-based approach that we used, the channel is held in a static conformation, allowing for an unambiguous calculation of drug-channel affinity for discrete channel conformations.

### Computational ion channel structure to function relationship

In this study we have attempted to make a novel link between ion channel structure and function. We utilized atomic scale predictions to inform rate constants for constructing computational channel-scale kinetic models for dofetilide interaction with hERG channels. Potential mean force calculations from drug – channel binding molecular-structure level trajectories allowed for the calculation of dissociation constants *K*D for dofetilide interactions with hERG for the open state of the channel. These simulated data combined with predicted diffusion coefficients from the same atomistic MD runs allowed for drug “on” and “off” rates to discrete states to be introduced into the function scale Markov model of hERG.

### Connection between structure – activity relationship and proarrhythmia

Computational models of dofetilide interaction with the hERG receptor were integrated into virtual cardiac cell and tissue level models to **predict** emergent drug effects to promote elements of the **TRIaD**: *Triangulation, reverse* use dependence (increase in drug effects at ***slow*** heart rates), beat-to-beat *instability* of action potential duration, temporal *and* spatial action potential duration *dispersion* – proarrhythmia markers that emerge at cell and tissue scales.

The driving hypothesis underlying the goals of this study is that the proarrhythmic cellular manifestations of the TRIaD arise directly from the underlying kinetics of channel block. Identification of the specific State-dependent kinetics of drug blockade that give rise to components of the TRIaD is essential to define new standards for preclinical compounds that can be used to rule out compounds with these properties in early screening tests. Our sensitivity analysis (Figure 7) suggests that as a general principle, reducing the affinity of hERG blocking drugs to the inactivated state will reduce the propensity to arrhythmias linked to **TRIaD** mediated arrhythmia vulnerability parameters. The sensitivity analysis suggests that the model was highly robust to small perturbations in the model parameters.

The manifestation of the **TRIaD** parameters can be observed in the tissue level simulations designed to serve as *in silico* diagnostic indication of arrhythmia vulnerability. In Figures 4 and 5, the model predictions show the emergence of arrhythmia triggers in the presence of low levels of applied electrical instability and dofetilide, both in the absence and in the presence of extra stimuli. The increase in instability and triangulation of the action potentials make cells “pre-treated” with dofetilide extremely vulnerable to small random spatial noise. It is the net effect of spontaneous depolarization that causes increased spatial and temporal action potential duration dispersion, combined with the profound reverse use-dependence of dofetilide that results in the extrasystolic induction of a reentrant wave following a pause in the presence of dofetilide as shown in Figure 5. Future studies must specifically test this concept in order to prove the mechanism and ultimately improve preclinical drug screening approaches.

Although it has long been clear that the **TRIaD** linked parameters indicate arrhythmia vulnerability (9, 28, 29), it has not been clear which are the minimal and sufficient parameters to predict arrhythmia risk. In order to better understand the required parameters that should be tracked in a simulation or an experiment, we used random forest machine learning algorithm to evaluate the importance of each arrhythmia vulnerability parameter from the **TRIaD** *(APD TRI, bTb instability, T-wave area and RUD)* to determine the final target classification (control versus drug affected at different dofetilide doses). The Pearson’s correlation coefficient was first calculated to explore the linear dependence between the **TRIaD** parameters. Not too surprisingly, we found that all **TRIaD** parameters are highly correlated, which indicates that there is no need to track them all because they contain redundant information. Given this outcome, we employed a multiclass random forest classification machine learning algorithm using the correlated **TRIaD** parameters to determine the relative importance of each **TRIaD** parameter to correctly predict whether a simulated cell belongs to the drug free, or low, medium or high risk category (indicated by dose in this study). Other drugs can be compared to dofetilide outputs to indicate risk relative to the high-risk drug dofetilide. The beat-to-beat (bTb) *instability* parameter was shown to be most important feature in classifying the drug safety.

In this study, we have brought together model simulations at the atomistic level for hERG channel structure, dynamics and channel – drug interactions as well as simulations at the functional levels of the protein, cell and tissue. *The power of combining these scales in a predictive framework is that it has allowed a way to derive on and off rates of drugs from atomic scale simulations and to then use these values to inform and build functional level channel models.* These function scale drug-channel models were then integrated into cellular and tissue level model to reveal mechanistic links between structure-activity relationships of ion channel – drug systems with higherorder emergent electrical phenomena such as cardiac rhythm disturbances. Our approach can be expanded for varied genotypes and myriad risk factors, and to predict individual responses to drug therapy.

Ultimately, our approach represents a scalable framework with automation potential to interact with other developing technologies, including high-throughput electrophysiology measurements (63–68) (69–74), drug development *via* progress in synthetic biology (75), and even personalized medicine *via* drug screening in patients’ own induced pluripotent stem (iPS) cell-derived cardiomyocytes (76). All of these developing technologies are innovative but each of them can’t alone solve the fundamental problem – that the effects of multifaceted drug interactions are ***emergent***. These technologies ***in conjunction*** with the multiscale models that we will develop may form *an interactive multiscale modeling and simulation driven process that can ultimately be used in the regulatory process prior to drug approval, in academia for research, in industry for drug and disease screening, and for patient-oriented medicine in the clinic.*

