## Supplemental methods for "A computational pipeline to predict cardiotoxicity: From the atom to the rhythm"

**Supplementary Materials**

**Fig. S1.**

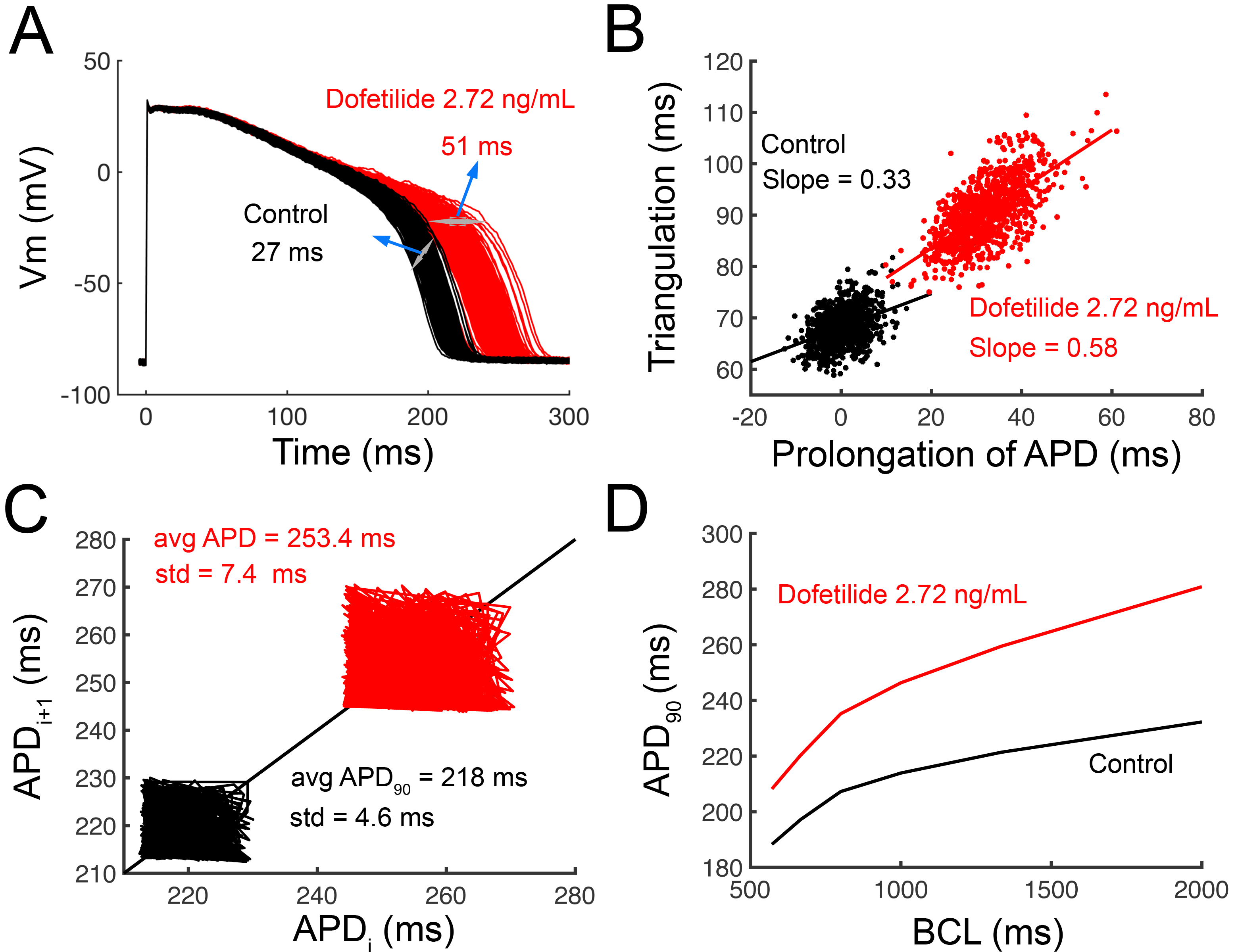

**Computational screen of arrhythmia vulnerability in rabbit model.** In panel A, ***temporal action potential duration* *dispersion*** was quantified in a cell population of 1000 individual simulated cardiac myocyte action potentials constructed by incorporating physiological noise (1, 2). *Dispersion* of APD was quantified as the difference between the maximum and minimum action potential duration. Dofetilide within the clinical dosing range has a clear effect to promote temporal action potiential duration variability in the presence of the drug (Control – 27 ms; Dofetilide 2.72 ng/mL = 51 ms). Panel B illustrates the effect of dofetilide to promote ***triangulation*** of the action potential as a function of APD prolongation. In the absence of drug, control cells had a slope = 0.33, while Dofetilide 2.72 ng/mL increased the slope = 0.58. Panel C shows Poincaré plots of sequential APD pairs indicating beat-to-beat ***instability*** following the application of small electrical perturbations in the absence of drug or with 2.72 ng/mL dofetilide. Instability was assessed by applying small amplitude inward currents randomly between -0.1 to -0.2 pA/pF for 50 ms over the course of the action potential plateau at a pacing cycle length = 1000 ms. Finally, in panel D, ***reverse use dependence*** induced by dofetilide was evaluated. The action potential adaptation curves were generated using APD_90_ values from human computational ventricular myocytes at steady-state at the indicated pacing frequencies. When dofetilide (red) was applied, there was a clear steepening of the APD adaptation curve compared to the baseline drug-free case (black).

1.	Sato D, Bers DM, & Shiferaw Y (2013) Formation of Spatially Discordant Alternans Due to Fluctuations and Diffusion of Calcium. *Plos One* 8(12).

2.	Tanskanen AJ & Alvarez LH (2007) Voltage noise influences action potential duration in cardiac myocytes. *Mathematical biosciences* 208(1):125-146.
